# Structural Model of Bacteriophage P22 Scaffolding Protein in a Procapsid by Magic-Angle Spinning NMR

**DOI:** 10.1101/2024.11.01.621488

**Authors:** Changmiao Guo, Richard D. Whitehead, Jochem Struppe, Gal Porat-Dahlerbruch, Alia Hassan, Angela M. Gronenborn, Andrei T. Alexandrescu, Carolyn M. Teschke, Tatyana Polenova

## Abstract

Icosahedral dsDNA viruses such as the tailed bacteriophages and herpesviruses have a conserved pathway to virion assembly that is initiated from a scaffolding protein driven procapsid formation. The dsDNA is actively packaged into procapsids, which undergo complex maturation reactions to form infectious virions. In bacteriophage P22, scaffolding protein (SP) directs the assembly of coat proteins into procapsids that have a T=7 icosahedral arrangement, en route to the formation of the mature P22 capsid. Other than the C-terminal helix-turn-helix involved in interaction with coat protein, the structure of the P22 303 amino acid scaffolding protein within the procapsid is not understood. Here, we present a structural model of P22 scaffolding protein encapsulated within the 23 MDa procapsid determined by magic angle spinning NMR spectroscopy. We took advantage of the 10-fold sensitivity gains afforded by the novel CPMAS CryoProbe to establish the secondary structure of P22 scaffolding protein and employed ^19^F MAS NMR experiments to probe its oligomeric state in the procapsid. Our results indicate that the scaffolding protein has both α-helical and disordered segments and forms a trimer of dimers when bound to the procapsid lattice. This work provides the first structural information for P22 SP beyond the C-terminal helix-turn-helix and demonstrates the power of MAS NMR to understand higher-order viral protein assemblies involving structural components that are inaccessible to other structural biology techniques.

## INTRODUCTION

Icosahedral viruses generally assemble by two basic mechanisms. 1) In a spontaneous reaction, the coat proteins (CPs) co-assemble around the genome. Viruses that assemble using this mechanism include polyoma viruses, HIV, hepatitis B, and some bacteriophages like MS2. 2) An immature, precursor icosahedral procapsid (PC) can be assembled into which the genome is actively packaged. Assembly of CPs into immature precursor capsids, or procapsids, is catalyzed by scaffolding proteins^1–4^. This mechanism is used by the tailed phages, herpesviruses, and adenoviruses. Scaffolding proteins (SP) directs assembly of coat proteins and also the incorporation of the portal protein dodecamer, and the internal ejection proteins that exit via the portal upon infection^5,6^.

Scaffolding proteins (SPs) of dsDNA viruses are similar in both structure and function. The coat-binding domain of P22 SP has a C-terminal helix-turn-helix (HTH) structure^7,8^. An α-helical region in the C-terminus of herpesvirus SP has also been identified as responsible for interaction with the capsid protein^9^. Helical and HTH motifs from SPs have also been found to direct assembly of phages SPP1, ϕ29 and 80α^10–13^. In addition, viral SPs are generally considered to have intrinsic disorder^14,15^, which is the reason that only two full-length SP structures from phages φ29^16^ and φBB1^17^ have been solved. Other than the C-terminal HTH, the structure of the P22 SP is unknown^18,19^.

Here, we report a structural model of the P22 scaffolding protein in the context of a 23-MDa large procapsid using magic angle spinning (MAS) NMR. We established the secondary structure of P22 SP using 2D and 3D ^13^C-detected experiments and probed the oligomeric state of SP in the procapsid by ^19^F MAS NMR. Our results indicate that the scaffolding protein has both α-helical and disordered segments and forms a trimer of dimers when bound to the procapsid lattice. Our work highlights the structural heterogeneity of the P22 scaffolding protein in the procapsid. We propose these techniques will be broadly applicable to other viral capsid and macromolecular assemblies that include proteins that are intrinsically disordered in solution.

## RESULTS AND DISCUSSION

### Local structure and conformational heterogeneity of the scaffolding protein in the P22 procapsid

2D ^13^C-^13^C cross-polarization (CP) based CORD experiment was performed to detect the rigid segments of SP. The CP-CORD spectrum contains both overlapping broad peaks and well-resolved resonances with strong intensities and narrow linewidths (Fig. 1b). Thus SP within the P22 procapsid has both structurally ordered rigid regions as well as statically disordered regions exhibiting conformational heterogeneity. A direct polarization (DP) based 2D DARR experiment was also conducted to see if SP possesses dynamic regions undergoing local motions on the milli-to microsecond timescales. The DP-DARR spectrum shown in Fig. 1c reveals additional strong cross peaks corresponding to Ala, Ile, Thr and Ser residues indicating that these are located in mobile regions of SP. Similarly, several correlations involving aromatic sidechains that are absent in the CP-based spectrum, are detected only in the 2D DP-DARR spectrum, indicating that these aromatic residues are dynamic. By residue type count and spin system identification we ascertained that more than 50% of the SP residues are detected in the 2D ^13^C-^13^C correlation spectra alone.

**Fig. 1.**
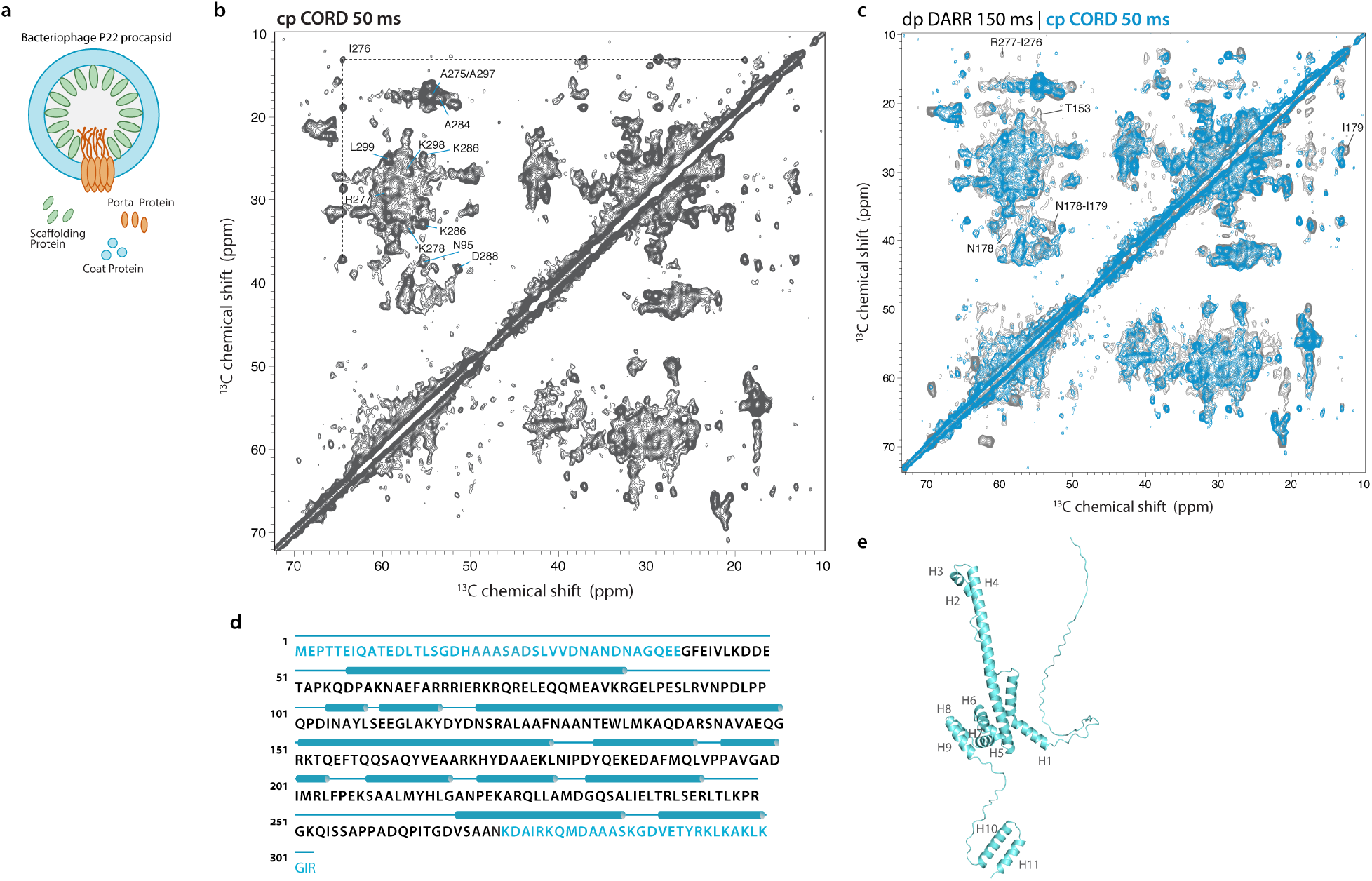
**a** Schematic representation of bacteriophage P22 procapsid assembly. **b** 2D ^13^C-^13^C cross-polarization (CP)-based CORD MAS NMR spectrum of U-[^13^C,^15^N]-labeled P22 SP assembled in PC; the mixing time was 50 ms. **c** 2D direct-polarization-based DARR spectrum (gray) mixing overlaid with 2D CP-CORD spectrum (blue); the mixing time was 150 ms. The spectra were acquired at 20.0 T; the MAS frequency was 14 kHz. **d** Sequence and secondary structure of the P22 scaffolding protein. The N- and C-terminal domains of SP are colored in cyan. The cylinders represent α-helices. **e** AlphaFold 3 structural model of the P22 SP monomer.

The sequence-based prediction and an AlphaFold 3 model of the SP monomer both indicate that P22 scaffolding protein has predominantly α-helical structure, linked by a significant proportion of disordered segments (Fig. 1 d,e). These structural features are consistent with the MAS NMR experimental ^13^C^α^ and ^13^C^β^ chemical shift distributions determined from the ^13^C^α^-^13^C^β^ correlations in the CP-CORD and DP-DARR spectra. Specifically, chemical shifts for Ala residues, highly abundant in the SP, indicate that most of the Ala residues possess α-helical secondary structure, with only a few comprising random coil, and none located in β-sheets. Similarly, most of the Thr, Ser and Ile residues have α-helical structure as well (Fig. 1b).

### P22 SP has a combination of rigid and dynamic regions

To gain information into the dynamic regions of the SP structure, we conducted additional scalar- and dipolar-based MAS NMR experiments. To reduce spectral complexity, we prepared the procapsid assembly using deuterated U-[^13^C,^15^N]-SP with 100% amide-proton back-exchanged and employed ^1^H-detected fast MAS NMR experiments, ^13^C-^1^H HETCOR, ^15^N-^1^H HETCOR and INEPT. A combination of rigid and highly dynamic segments of SP in the procapsid were detected in these experiments. ^1^H-detected MAS NMR ^15^N-^1^H HETCOR and INEPT spectra corroborate that SP in the procapsid is conformationally heterogeneous, as is evident from severe peak overlap in the ^1^H dimension despite the narrow ^1^H linewidth (Fig. 2a). Interestingly, the CP-based ^15^N-^1^H HETCOR spectra reveal several broad intense peaks that correspond to the HTH residues of the C-terminal domain (CTD), indicating that the HTH as the capsid-binding domain is rigid and conformationally heterogeneous. Moreover, there are numerous peaks possessing narrower linewidths and relatively low intensities, which correspond to the residues in the molten globule domain.

**Fig. 2.**
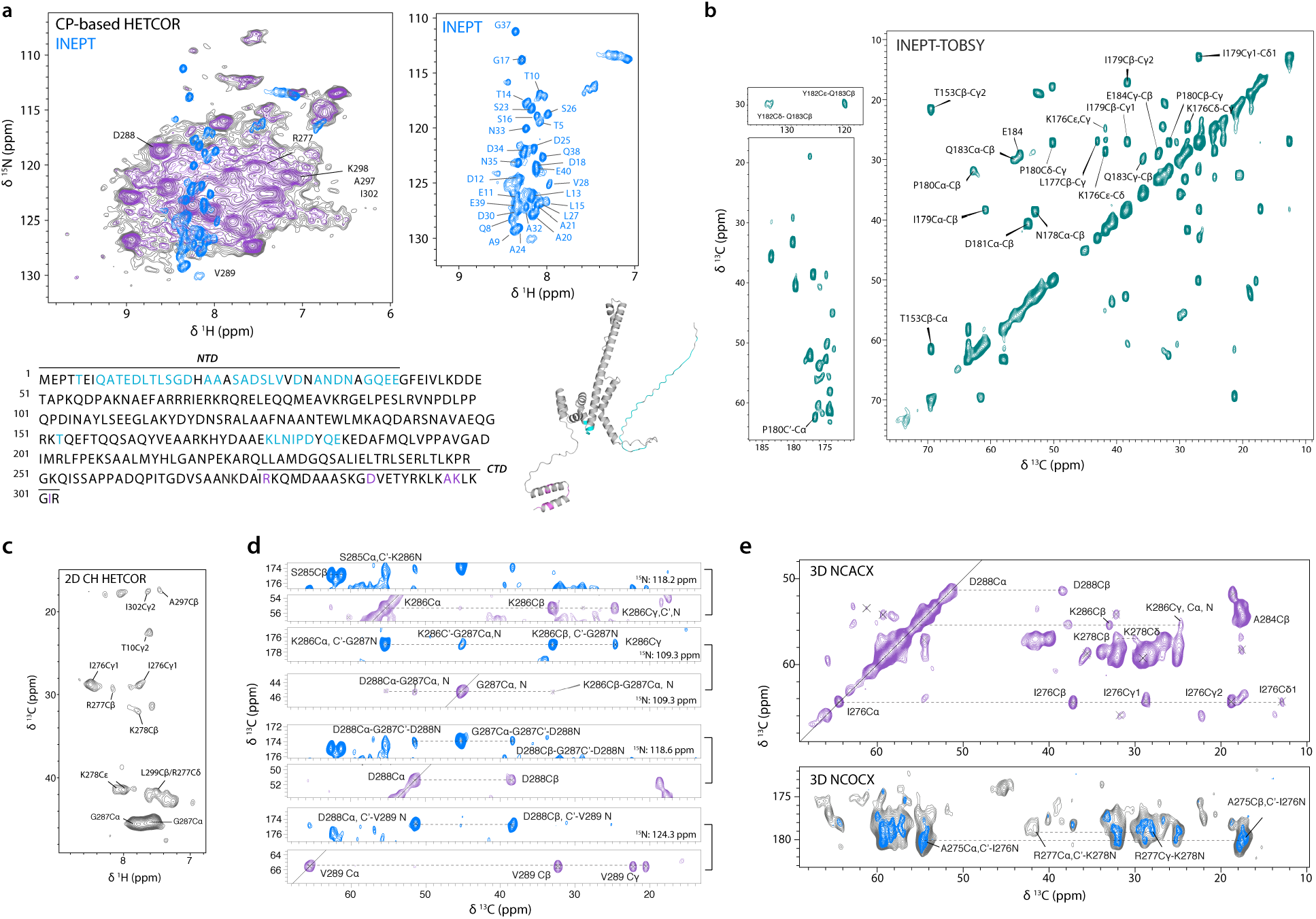
**a** 2D dipolar-based ^15^N-^1^H HETCOR and scalar-based INEPT MAS NMR spectra of deuterated U-[^13^C,^15^N]-labeled P22 SP assembled in PC. The CP-based HETCOR spectrum with resolution enhancement processing is shown in purple. Rigid and highly dynamic residues in SP are mapped to the SP sequence and model, colored in purple and cyan, respectively. **b**,**c** 2D ^13^C-^13^C scalar-based INEPT-TOBSY and 2D dipolar-based ^13^C-^1^H HETCOR spectrum of U-[^13^C,^15^N]-SP PC assembly. **d**,**e** Backbone resonance assignments for the S285-V289 segment in P22 SP assembly (**d**), using the 3D ^13^C-detected NCACX and NCOCX spectra (**e**) acquired with the CPMAS CryoProbe.

**Fig. 3.**
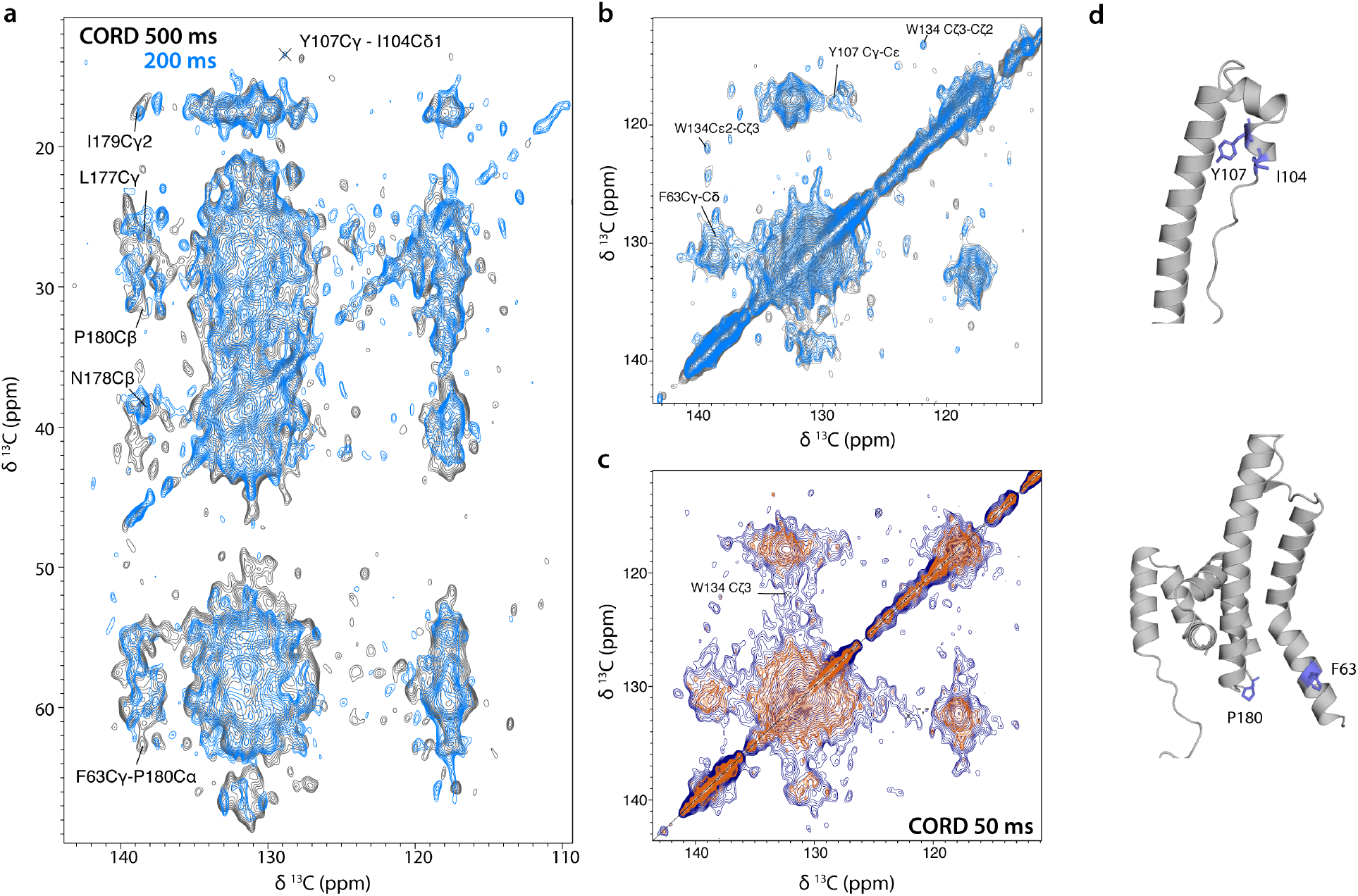
**a**,**b** 2D CORD spectrum of P22 SP assembled in PC acquired with mixing times of 200 ms (blue) and 500 ms (gray). The aliphatic-aromatic and aromatic-aromatic regions are shown in **a** and **b**, respectively **c** Aromatic contacts in the 2D CORD spectrum with 50 ms. **d** Aromatic-based restraints mapped on the SP model.

Highly dynamic SP residues were probed by scalar-based correlation experiments. Scalar-based 2D ^15^N-^1^H INEPT spectrum primarily reveals the amide correlations of the SP N-terminal domain (NTD). The NTD residues are altogether absent in the CP-based spectrum, consistent with NTD being highly flexible in the procapsid (Fig. 2a). Interestingly, the K176-E184 stretch of residues is clearly detected in the 2D ^13^C-^13^C scalar-based spectrum with through-bond TOBSY mixing (Fig. 2b), hence constituting an additional dynamic segment. This region is the linker between helix 4 and helix 5, and the NMR data indicate that, surprisingly, it has higher dynamics than the disordered segment between the molten globule domain and the CTD, as the latter was not detected by the INEPT-based experiments.

Of note is that MAS NMR characterizations of SP in the P22 procapsid are challenged by the low sensitivity and high complexity of this assembly, as the proportion of isotopically labeled SP is less than 10% and the procapsid is a multicomponent heterogeneous assembly. To overcome the low sensitivity, we employed the CPMAS Cryo-Probe that yielded 10-fold compounded sensitivity gains in 1D experiments^20^ and enabled acquisition of numerous 3D ^13^C-detected heteronuclear correlation spectra, not feasible with conventional MAS probes. Using a combination of these 3D spectra and 2D CORD spectra, we performed partial site-specific resonance assignments of P22 SP, with complete backbone assignments accomplished so far for stretches of residues A275-K278, A283-V289 and A297-L299 in the CTD (Fig.2 d,e). These CPMAS CryoProbe spectra were also used to corroborate that the aliphatic carbon-to-amide proton contacts present in the 2D ^13^C-^1^H HETCOR spectrum mostly correspond to the CTD residues. Furthermore, we ascertained that multiple peaks are observed for several residues such as G287, K278 and I276 arise from the conformational heterogeneity of the CTD in different SP chains bound to the capsid lattice (Fig. 2c), consistent with the low overall symmetry of SP in the procapsid (see below).

### Long-range contacts

Numerous long-range restraints involving aromatic residues are detected in the 2D CORD spectra with mixing times above 200 ms. These include a number of aliphatic-aromatic and aromatic-aromatic contacts. These long mixing times CORD spectra were also very helpful in making additional residue-specific assignments, based on the identifications of the spin systems and residue sequential connectivity patterns.

### Oligomeric state of SP in the P22 procapsid

To ascertain the oligomerization of SP in the P22 procapsid, we used ^19^F as a reporter and acquired ^19^F MAS NMR spectra of 5F-Trp-SP PC assemblies. Specifically, two procapsid samples were prepared, where 5F-Trp was incorporated into W-134 (WT SP) and W-10 (T10W/W134Y SP variant) (Fig. 4a). In the WT SP, W134 is located in the molten globule domain, whereas the T10W/W134Y variant contains W10 in the NTD. The 1D ^19^F MAS NMR spectrum of 5F-Trp substituted WT SP shown in Fig. 4b reveals multiple overlapping ^19^F resonances that span an overall chemical shift range of ca. 6 ppm. In contrast, the 5F-Trp T10W/W134Y SP mutant yields a single intense ^19^F peak with a linewidth of ∼1-2 ppm, although there are more than one ^19^F subspecies. Interestingly, the individual ^19^F resonances in both samples are relatively narrow, with the estimated homogeneous linewidths of 1 ppm determined on the basis of the DANTE-based selective inversion experiment. Indeed, the DANTE selective-inversion spectra of 5F-Trp WT SP indicate that the broad peak consists of several distinct ^19^F species. Taken together, these experimental results indicate that P22 SP forms higher-order oligomers in the procapsid and the segment encompassing residue W134 is involved in the SP oligomerization inside the PC lattice. On the other hand, the W10 of the NTD is located in a less heterogeneous structural environment compared to W134, albeit structural variations are still present. The dramatic difference between the ^19^F spectra of WT and mutant SP highlights the structural heterogeneity of the molten globule domain.

**Fig. 4.**
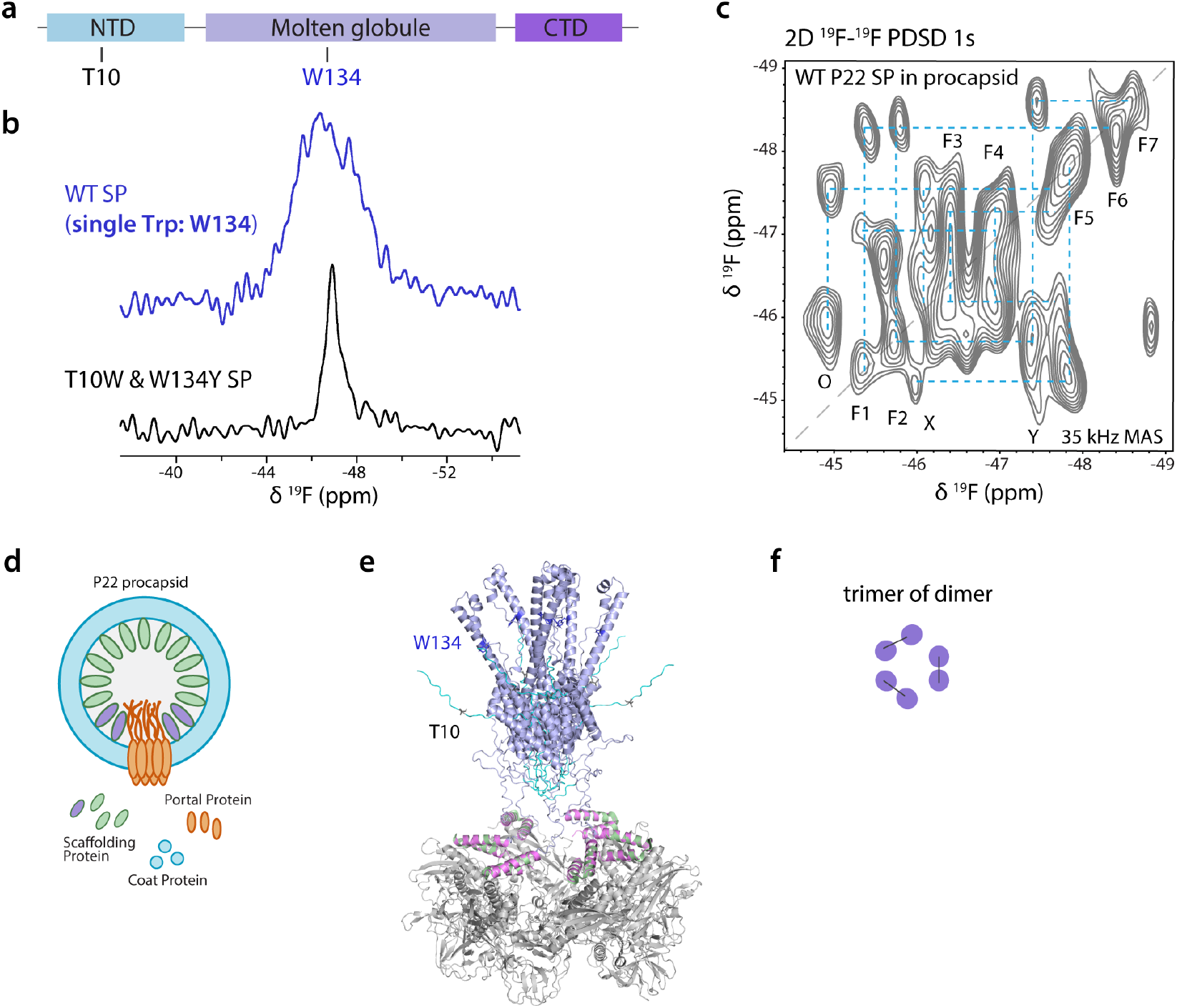
**a** Domain organization of P22 SP. **b** 1D ^19^F MAS NMR spectrum of 5F-Trp WT SP (purple) and T10W/W134Y SP variant (black) assembled in PC. **c** 2D ^19^F-^19^F dipolar-based spin diffusion (SD) MAS NMR spectrum of 5F-Trp WT SP PC assembly. **d** Schematic illustration of the P22 PC assembly to show the structural variations of SP pairs near the portal vertex and interacting with the capsid lattice. **e** Structural model of the P22 SP oligomer in PC. ESMFold (ref) model of SP and cryo-EM density map of SP CTD and coat proteins (PDB 8I1V^19^) were used. **f** Schematics of the possible arrangements for the trimer of dimers of P22 SP in the PC.

To understand the oligomeric state of the SP, we recorded a 2D ^19^F-^19^F dipolar-based spin diffusion (SD) spectrum on the 5F-Trp WT SP assembled in PC. The SD experiment reports on the spatial proximity of the dipolar-coupled ^19^F species belonging to the different 5F-Trp residues in each SP chain. Remarkably, the SD spectrum shown in Fig. 4c. reveals multiple strong cross peaks indicating that the 5F-Trp residues in the different local environments are separated by less than 10 Å. Moreover, the cross peak pattern is consistent with the overall symmetry being either a dimer-of-trimers or trimer-of-dimers or at least two distinct types of hexamers. Based on the spectrum, both dimer-of-dimers and a tetramer arrangement can be clearly ruled out based on the following considerations. First, seven distinct diagonal peaks and fifteen ^19^F-^19^F cross peaks were detected. Second, there are correlations between different cross peak pairs, suggesting that the SP forms higher-order symmetry elements, beyond dimers. Third, the 2D spectrum reveals more than 6 sets of ^19^F resonances, which indicates structural variations of SP oligomers. Lastly, there is no evidence the SP ever forms trimers^21,22^; thus, we hypothesize SP forms a trimer of dimers inside PCs based on the spectrum. These results are consistent with the previously reported structures of the P22 procapsid assembly^23^. Chiu and coworkers have determined the asymmetric structure of icosahedral P22 procapsid by cryo-EM and reported a symmetry mismatch for the scaffolding protein that interacts with the portal^18^. They found that each hexamer of the capsid associates with two scaffolding proteins at the portal vertex, and there are variations in the orientations of each SP dimer near the portal. Conway and coworkers explained the similar symmetry mismatch in the bacteriophage HK97 procapsid by flexible accommodation of a 12-fold pore at a 5-fold viral capsid vertex^24^. Therefore, the structural variations and heterogeneity of the scaffolding protein revealed by ^19^F MAS NMR may arise from the SP pairs that interact with the portal as well as the hexamer and pentamer of the capsid lattice^18,19^.

## Discussion

Scaffolding proteins of icosahedral dsDNA viruses and bacteriophages mediate the assembly of coat proteins into procapsid. Atomic-level structures of SP within the procapsid are critical for understanding how SP drives the assembly of procapsid as well as the SP release and genome packaging. P22 SP represents a challenging system for structural studies as it forms oligomers with conformational heterogeneities and contains a significant proportion of disordered and dynamic regions in the procapsid. Therefore, visualization of the EM density of full-length SP encapsulated in the shell is challenging. As SP has a combination of ordered and disordered regions, it remains difficult for artificial intelligence (AI)-based structure prediction.

We demonstrated that MAS NMR provides atomic-level information for the secondary and tertiary structure, conformational heterogeneities, and structural dynamics of scaffolding proteins. Our results show that high-resolution NMR spectra can be obtained despite of the complexity of the P22 procapsid. Using different MAS NMR experimental schemes, structurally ordered regions as well as the statically and dynamically disordered regions of P22 SP are detected. This work showcases that a combination of the novel CPMAS CryoProbe technology and ^19^F-as well as ^1^H-detected fast MAS NMR provide sensitivity and resolution gains that allow atomic-level structural analysis of SP in viral procapsid. Our work demonstrates that MAS NMR is a powerful tool for understanding higher order viral assemblies involving structural components and intrinsically disordered regions that are inaccessible to other structural biology techniques.

## ACKNOWLEDGMENTS

This work was supported by the National Institutes of Health (NIH grant R01GM076661). We acknowledge the support of the National Science Foundation (NSF grant CHE-0959496) for the acquisition of the 850 MHz NMR spectrometer at the University of Delaware. We thank Dr. Chunting Zhang for her involvement in the NMR experiments at the initial stage of the project. We acknowledge the support by the National Institutes of Health NIH (Grants P50AI1504817 and U54AI170791, NMR Core) that enabled the technology development required for MAS NMR structural characterization of complex viral protein assemblies.

## AUTHOR CONTRIBUTIONS

C.M.T. and T.P. conceived the project. C.G. performed the MAS NMR experiments, processed all the NMR data, performed the data analysis and resonance assignments. R.D.W. prepared the samples and conducted biochemical characterizations. J.S., T.P., A.H., and A.M.G. performed the CPMAS CryoProbe-based experiments. G.P.D. and C.G. performed the NMR experiments on deuterated samples. G.P.D. processed the 3D spectra and did partial backbone assignments on the CTD. C.G., R.D.W., T.P., and C.M.T. took the lead in writing the manuscript with input from all authors. All authors approved the final version of the manuscript.

## MATERIALS AND METHODS

### Chemicals

Ammonium chloride (^15^N, 99%), U-^13^C_6_ glucose, U-[^13^C_6_, D_7_]-glucose and D_2_O (99.8 % purity) were purchased from Cambridge Laboratories, Inc.

### Protein expression and purification

A pET15b vector containing the 6x-His tagged full length scaffolding protein (SP) was transformed into BL21 (DE3) cells. For expression of uniform isotopically labeled SP, cells were grown in M9 minimal media supplemented with 1 g/L ^15^NH_4_Cl or 3 g/L U-^13^C glucose (Cambridge Isotope Laboratories, Cambridge, MA). After inoculation of M9 media using a starter culture, the cells were grown to an OD_600_ of 0.5 at 37°C before induction of SP with 1 mM IPTG (Gold Biotechnology, St. Louis, MO). For preparation of 5F-Trp-SP, 20 mg/L of 5-flourindole in 1 mL 70 % ethanol (70/30 ethanol/water v/v) was added to M9 media 1 hour prior to induction. After induction, cells were grown for 16 hours at 18°C and harvested via centrifugation at 7,800 *g* for 15 minutes. The cells were resuspended in 20 mM HEPES buffer, 20 mM Imidazole, pH 7.4.

Cells were lysed via sonication (Misonix, Farmingdale, NY) at an amplitude of 37 for a total time of 3 min (15 s pulse on and 30 s pulse off). Cell debris were pelleted by centrifugation at 31,920 *g* for 15 min. The supernatant was transferred to a 75°C water bath for 10 minutes to denature the bulk of the *E. Coli* proteins before chilling on ice for 10 minutes to allow for SP refolding. The protein solution was spun at 31,920 *g* for 10 minutes to pellet the heat-denatured proteins. The supernatant was loaded onto a Ni-NTA agarose affinity column (Qiagen, Hilden, Germany) and washed with 30 mL HEPES buffer (20 mM HEPES, 20 mM imidazole). The P22 SP was eluted with 20 mM HEPES, 250 mM imidazole using a 0-100% linear gradient. Fractions containing His_6_-SP were identified by SDS-PAGE. SP was subsequently dialyzed into water and lyophilized both for concentrating and long-term storage.

### Encapsulation of SP into P22 procapsids

Procapsid shells lacking SP were generated as previously described^25^. Isotopically labeled SP was incubated with natural abundant empty P22 procapsid shells at a ratio of 100:1 SP:shell to ensure tight binding of SP to coat protein. The PC shell was restuffed with scaffolding protein in 20 mM sodium phosphate, 50 mM NaCl, pH 7.5. The mixture was incubated at 25°C for 1 hour before sedimentation of the complex at 175,000 *g* for 20 minutes using a RP80AT rotor (Thermo Scientific, Waltham MA), in a Sorvall RC M120EX ultracentrifuge (Thermo Scientific, Waltham MA). The pellet was resuspended in a small volume of sodium phosphate buffer (20 mM NaPO_4_, 50 mM NaCl, 2 mM EDTA, 0.2% w/v sodium azide, pH 6.5) on a reciprocal shaker (Eberbach, Ann Arbor, MI) at 180 osc/min for ∼20 hours at 4°C.

P22 SP PC assemblies were pelleted down by ultracentrifugation at 200,000 *g* for 45 minutes at 4 °C. 3.8 mg and 45.4 mg hydrated pellets were packed into 1.3 mm and 3.2 mm Bruker MAS NMR rotors, respectively; 92 mg of PC complex was packed into a 3.2 mm Cryo-Probe MAS rotor.

### MAS NMR spectroscopy

The ^1^H, ^13^C and ^19^F-detected MAS NMR experiments were performed on 20.0 T (^1^H Larmor frequency of 850.4 MHz) Bruker Avance III NMR spectrometer. 2D and 3D ^13^C-detected MAS NMR spectra were acquired on uniformly ^13^C,^15^N-labeled SP in assemblies with coat proteins at natural abundance using a 3.2 mm EFree HXY probe. Typical 90° pulses were 2.65 μs for ^1^H, 4.3 μs for ^13^C and 4.4 μs for ^15^N. Homonuclear 2D CP-CORD and DP-DARR spectra of protonated P22 PC were acquired at a MAS frequency of 14 kHz. SPINAL64 scheme was used for ^1^H decoupling at a radio frequency (rf) field of 90 kHz during ^13^C and ^15^N evolutions.

^1^H-detected spectra of deuterated U-[^13^C,^15^N]-P22 SP in the procapsid were acquired at a MAS frequency of 60 kHz using a 1.3 mm HXY MAS probe. ^1^H decoupling was applied at a rf field of 15 kHz using the swept-frequency two-pulse phase modulation (SW_f_-TPPM) scheme. WALTZ-16^25^ scheme was applied on ^13^C and ^15^N at rf field of 15 kHz during ^1^H evolution.

The Cryo-Probe-based experiments were performed on 14.1 T (^1^H Larmor frequency of 600.3 MHz) Bruker Avance NEO NMR spectrometer equipped with a 3.2 mm HCN CPMAS CryoProbe. Heteronuclear N-CA and N-CO transfers were established using SPECIFIC-CP. Experimental parameters of all 2D and 3D ^13^C- and ^15^N-detected experiments are:

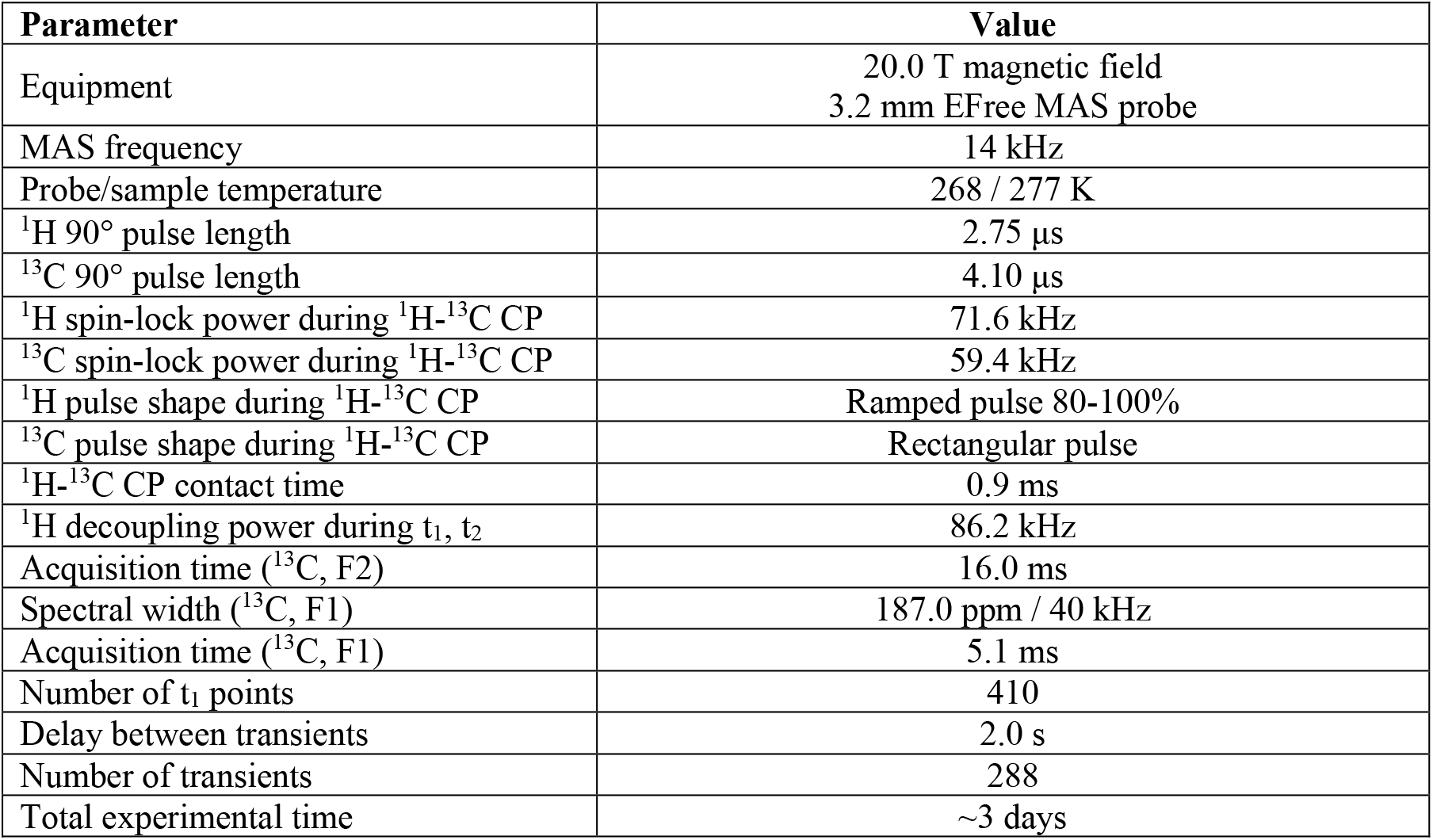

^19^F-detected experiments were performed using a 1.9 mm HX MAS probe. The ^1^H channel was tuned to ^19^F Larmor frequency of 800.1 MHz. All ^19^F-detected spectra were acquired on 5F-Trp, U-[^15^N]-SP PC assembly at a MAS frequency of 35 kHz. ^19^F pulse length was 1.85 μs and inter-scan delay was 5 s. 1D ^19^F experiments were acquired through triple-pulse excitation scheme to suppress probe background (69). 2D ^19^F-^19^F PDSD spectrum was acquired with a single-pulse excitation and spin diffusion of 1 s. A total of 24 points were acquired in t_1_ with 2,304 transients per increment. The acquisition time in t_1_ was 1 ms with a spectral width of 14.6 ppm.

### NMR data processing and analysis

^1^H and ^13^C-detected spectra were processed using TopSpin 4.0.2. Heteronuclear 3D NCACX and NCOCX spectra were processed with NMRPipe on NMRbox. 2D ^13^C-^13^C spectra were processed with apodizations Gaussian function of 40 Hz for sensitivity enhancement or 60-degree shifted sine bell function for resolution enhancement.

All NMR spectra were analyzed using CcpNmr analysis version 2.4.0.

### Structure prediction and modeling

Structural model of P22 SP single chain was predicted by AlphaFold 3^26^ and ESMFold^27^. The model of SP within the procapsid was generated by docking the ESMFold structure of SP monomer to the cryo-EM structure of P22 procapsid (PDB entry 8I1V^19^).

